# Natural history of nonhuman primates after conjunctival exposure to Ebola virus

**DOI:** 10.1101/2021.12.29.474476

**Authors:** Robert W. Cross, Abhishek N. Prasad, Courtney B. Woolsey, Krystle N. Agans, Viktoriya Borisevich, Natalie S. Dobias, Jason E. Comer, Daniel J. Deer, Joan B. Geisbert, Angela L. Rasmussen, W. Ian Lipkin, Karla A. Fenton, Thomas W. Geisbert

**Author notes:** **Correspondence:** Thomas W. Geisbert, Ph.D., University of Texas Medical Branch, Galveston National Laboratory, 301 University Blvd, Galveston, TX 77550-0610.

## Abstract

Transmission of Ebola virus (EBOV) primarily occurs via contact exposure of mucosal surfaces with infected body fluids. Historically, nonhuman primate (NHP) challenge studies have employed intramuscular or small particle aerosol exposure, which are uniformly lethal routes of infection, but mimic worst-case scenarios such as a needlestick. When exposed by more likely routes of natural infection, limited NHP studies have shown delayed onset of disease and reduced mortality. Here we performed a series of systematic natural history studies in cynomolgus macaques with a range of conjunctival exposure doses. Challenge with 10,000 plaque forming units (PFU) of EBOV was uniformly lethal, whereas 5/6 subjects survived low and moderate dose challenges (100 or 500 PFU). Conjunctival challenge resulted in a protracted time-to death. Asymptomatic disease was observed in survivors with limited detection of EBOV replication. Inconsistent seropositivity in survivors may suggest physical or natural immunological barriers are sufficient to prevent widespread viral dissemination.

## Introduction

Filoviruses, including Ebola virus (EBOV) and Marburg virus (MARV), are responsible for periodic outbreaks of viral hemorrhagic fever with case fatality rates of up to 90% (Feldmann 2013). Historically, filoviruses have caused relatively small outbreaks ranging from one case to a few hundred cases with limited geographic spread (Feldmann 2013). The 2013-2016 West African epidemic of EBOV (strain Makona) represented a departure from this pattern, with over 28,000 cases and 11,000 deaths (Coltart, Lindsey et al. 2017). More recently, the resolved 2018-2020 outbreak of EBOV (strain Ituri) (McMullan, Flint et al. 2019) in the Democratic Republic of the Congo (DRC) resulted in at least 3,481 cases and 2,299 deaths (WHO 2020). The increased scale, duration, and geographic footprint of these outbreaks emphasize the gaps in understanding the variety of modes of human-to-human transmission of filoviruses. While fractured healthcare systems, socio-political unrest, and security issues are known to contribute to spread of the virus, clarification is needed in the specific role different routes of infection play in the kinetics of virus transmission and disease progression, particularly in mild, asymptomatic, or recrudescent cases.

Irrespective of scale, filovirus outbreaks in humans have frequently been traced back to a single episode of zoonotic transmission, after which human-to-human transmission drives the remainder of the outbreak (Georges, Leroy et al. 1999, Leroy, Rouquet et al. 2004, Marí Saéz, Weiss et al. 2015). Instances of multiple zoonotic introductions within a single outbreak have also been reported (Swanepoel, Smit et al. 2007, Adjemian, Farnon et al. 2011). Egyptian rousette bats (*Rousettus aegyptiacus*) play a role in the natural maintenance of MARV (Towner, Amman et al. 2009, Amman, Carroll et al. 2012), and several species of frugivorous and insectivorous bats have been implicated, but not confirmed, as potential reservoir species for EBOV (Olival and Hayman 2014). In addition, secondary species, such as nonhuman primates (NHP) and duikers, may facilitate spillover of filoviruses into humans (Caron, Bourgarel et al. 2018). Intramuscular (i.m.) inoculation through the re-use of needles in a clinical setting fueled the spread of EBOV during the initial outbreak in Yambuku, DRC in 1976 (WHO 1978), but has not played a major role in subsequent outbreaks. Still, percutaneous/parenteral exposure via needlestick injury is an ever-present hazard to personnel in both research and clinical environments (Stone 2004, Gunther, Feldmann et al. 2011, Jacobs, Aarons et al. 2015, Rubinson 2015). The route of infection in cases with no documented needle use is less clear, but is presumed to involve mucosal surfaces (e.g., conjunctival, oral, nasal, or sexual exposure) (Vetter, Fischer et al. 2016). EBOV is found in a variety of bodily fluids including saliva, blood, stool, breast milk, and semen, rendering it highly contagious. Indeed, the primary risk factor for human-to-human transmission of filoviruses is mucosal contact with body fluids from infected persons, including sexual contact (Fischer and Wohl 2016).

A clear understanding of the natural history of different routes of infection is critical for understanding the epidemiology of pathogenic agents and for the development of medical countermeasures. This is particularly important in the case of filovirus disease, as most preclinical development and validation efforts of vaccines and therapeutics in animal models, such as NHPs, have demonstrated protection against i.m. injection or small particle aerosol rather than mucosal exposures. Thus, it is largely unknown how the protective efficacy of these countermeasures may be impacted by different routes of filovirus infection. While implementation of the US Food and Drug Administration-approved EBOV vaccine ERVEBO has had clear benefit in the management of EBOV outbreaks (Tomori and Kolawole 2021), there is still much to be learned regarding what impacts mucosal exposure have on medical countermeasure efficacy.

NHPs have historically been used as the animal model of choice against filoviruses as they recapitulate the most salient features of fatal human EBOV disease (EVD) (Geisbert, Strong et al. 2015). Previous work has demonstrated that low doses (0.01-50 PFU) of EBOV or MARV delivered by i.m. injection cause lethal disease in NHPs (Sullivan, Geisbert et al. 2003, Alfson, Avena et al. 2015, Alfson, Avena et al. 2018, Woolsey, Geisbert et al. 2018). Limited data exist characterizing the required doses needed to cause an infection. Moreover, the natural history of mucosal infection by filoviruses in NHPs is largely unexplored. Several studies have examined small particle aerosol challenge to understand disease processes and to evaluate medical countermeasures in the context of an intentional filovirus release. (Johnson, Jaax et al. 1995, Geisbert, Daddario-Dicaprio et al. 2008, Alves, Glynn et al. 2010, Reed, Lackemeyer et al. 2011, Zumbrun, Bloomfield et al. 2012, Twenhafel, Mattix et al. 2013, Ewers, Pratt et al. 2016). A recent study characterized an EBOV intranasal challenge model using a large particle generating atomizer, a model more reflective of person-to-person airborne droplet transmission (Alfson, Avena et al. 2017). In this study, disease progression was delayed compared to i.m. challenge with the large particle but not small particle aerosol challenge. These results suggest the virus faces additional physical and/or immunological barriers to infection via inhalational or mucosal routes tied to droplet size. Two studies have examined a range of infectious doses in both rhesus and cynomolgus macaques by the oral and conjunctival routes (Jaax, Davis et al. 1996, Mire, Geisbert et al. 2016); however, the outcomes were limited by the narrow scope and small group sizes in both studies. The work here represents a systematic extension of previous work in cynomolgus monkeys. Here, we add to the body of knowledge concerning the natural history of lethal disease and survival to three different doses of conjunctival exposure to EBOV using the 2013-2016 epidemic Makona variant.

## Results

### Experimental infection of cynomolgus macaques with EBOV via the conjunctival route

We exposed three groups (n=6 animals/group) of healthy, adult cynomolgus macaques with a target dose of 100, 500, or 10,000 PFU of EBOV via the conjunctiva. In both lower dose cohorts, surviving animals (5/6 per group) showed minimal clinical signs of disease, with decreased food intake or brief anorexia being the only outward symptom (**Table S1**). Only animals that progressed to fatal disease exhibited sustained fever followed by progressive hypothermia (**Figure S1**). Conversely, each of the animals that developed fatal disease in these two cohorts developed classical clinical signs of Ebola virus disease (EVD), including anorexia, recumbency, and petechial rash, while fever was only present in one of the animals. Both animals met euthanasia criteria 11 days post-infection (dpi) (**Figure 1A**). In contrast to the two lower dose groups, all the animals in the high challenge dose group developed fatal EVD, with a mean time to death (MTD) of 12.33 ± 3.25 dpi. The disease course in 3/6 animals was extended to 13-17 days, with the first clinical signs of illness appearing 6-11 dpi, while the remaining three animals met euthanasia criteria on days 9 or 10. There was a significant difference in survival curves between the 100 and 10,000 PFU challenge cohorts (p = 0.0049, Mantel-Cox log rank test) and 500 and 10,000 PFU cohorts (p = 0.0049, Mantel-Cox log rank test), but not between the 100 and 500 PFU cohorts (p > 0.99, Mantel-Cox log rank test) (**Figure 1A**).

**Figure 1:**
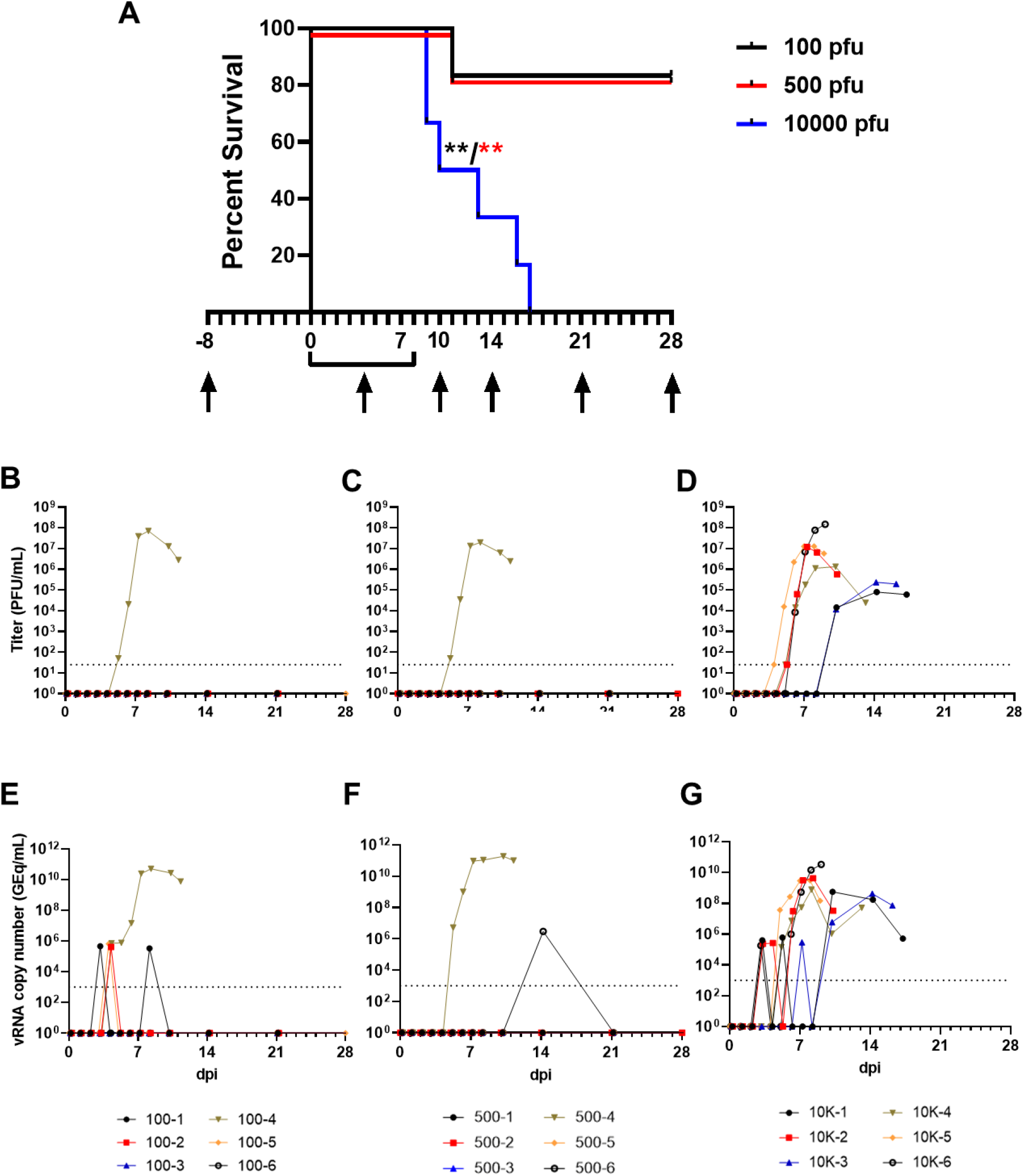
Survival analysis and determination of viral load in EBOV-challenged macaques. **(A)** Kaplan-Meier survival curves of cynomolgus macaques challenged with low (100-500 PFU) and high (10,000 PFU) doses of EBOV Makona. Arrows below x-axis denote scheduled sampling days. Viral load was determined by plaque titration of plasma (**B-D)** and RT-qPCR detection of EBOV vRNA **(E-G)** from whole blood.

Marked changes to blood coagulation parameters (e.g., increased prothrombin time (PT) and activated partial thromboplastin time (aPTT), decreased circulating fibrinogen). These findings are indicative of severe disruption to extrinsic and intrinsic coagulation pathways (i.e., acute disseminated intravascular coagulation and liver damage/failure was observed in all fatal cases regardless of challenge dose) (**Figure S2A-C**). A single surviving animal from the 500 PFU challenge cohort (subject 500-4) displayed similar deviation from baseline coagulation indices which resolved completely by 14 dpi (**Figure S2B**). Predictably, all fatal cases exhibited hallmark features documented in other reports of NHP infection with EBOV regardless of dose. Dramatic changes in leukocyte populations were observed in these subjects, as well as serum markers of hepatic and renal function and inflammation, compared to those that survived (**Table S1**). Notably, in fatal cases, there was a lack of association between the challenge dose and the severity of disruption from baseline hematological and metabolic parameters.

### Quantitation of EBOV vRNA and infectious virus load

We assessed levels of circulating infectious virus in plasma and viral RNA (vRNA) in whole blood by plaque assay and RT-qPCR, respectively. In both lower dose groups, infectious EBOV was only recovered in animals that developed fatal disease (**Figure 1B,C**) while all animals developed levels of circulating infectious virus in the high dose group (**Figure 1D**). In the two animals from the high dose cohort exhibiting the most protracted time-to-death (TTD) (euthanized at 16 and 17 dpi), detectable levels of circulating infectious EBOV and vRNA developed later and at lower titers than in the animals with a shorter disease course (**Figure 1D,G**). Circulating EBOV vRNA was detected transiently in 3/6 and 1/6 survivors from the 100 and 500 PFU challenge cohorts, respectively (**Figures 1E,F**). Tissues were collected at necropsy and assayed for the presence of EBOV vRNA and infectious virus. vRNA was primarily restricted to lymphoid tissue, liver, spleen, and lungs in surviving animals from the 100 PFU cohort (**Figure S3A**) but was also found in low abundance (~10^4^ GEq/g tissue) in the eye and transverse colon of one survivor (100-5). Similarly, EBOV vRNA was absent in most tissues from surviving animals in the 500 PFU challenge cohort; however, two animals (500-1 and 500-6) had detectable vRNA in the gonads, and vRNA was found in the eyes from three survivors (500-1, 500-3, 500-6) (**Figure S3C**). vRNA was present in similar quantities (~10^8^-10^10^ GEq/g tissue) in most or all tissues from all animals which succumbed, regardless of challenge dose (**Figure S3A,C,E**). Likewise, titers of infectious virus recovered from the tissues of animals which succumbed also did not appear to be dependent on the challenge dose (**Figure S3B,D,F**). Infectious virus was absent, or below the limit of detection, in the pancreas of 3/6 animals (10K-4, 10K-5, 10K-6) in the 10,000 PFU challenge cohort, but was recovered from all other animals that succumbed. Infectious virus was not recovered from any tissues assayed from surviving animals. There was no significant difference in the mean peak viral load, day of peak viremia, or day of earliest detection, whether measured by plaque assay (for fatal cases only) or RT-qPCR (for all animals with detectable vRNA), between the two lower dose groups and the high dose group (**Figure S4A,B**), although fatal cases had markedly higher levels of vRNA in both blood and tissues (**Figure S4B**).

### Pathology

Regardless of challenge dose, postmortem gross examination of animals which succumbed to lethal disease revealed lesions consistent with those observed in i.m. challenge models, including necrotizing hepatitis (**Fig 2B & C**) characterized as hepatic pallor with reticulation; splenomegaly; petechial rash on the limbs, face, and/or trunk (**Fig 2D**); and hemorrhagic interstitial pneumonia characterized as failure to completely collapse and multifocal reddening of the lungs (**Fig 2E**).

**Figure 2:**
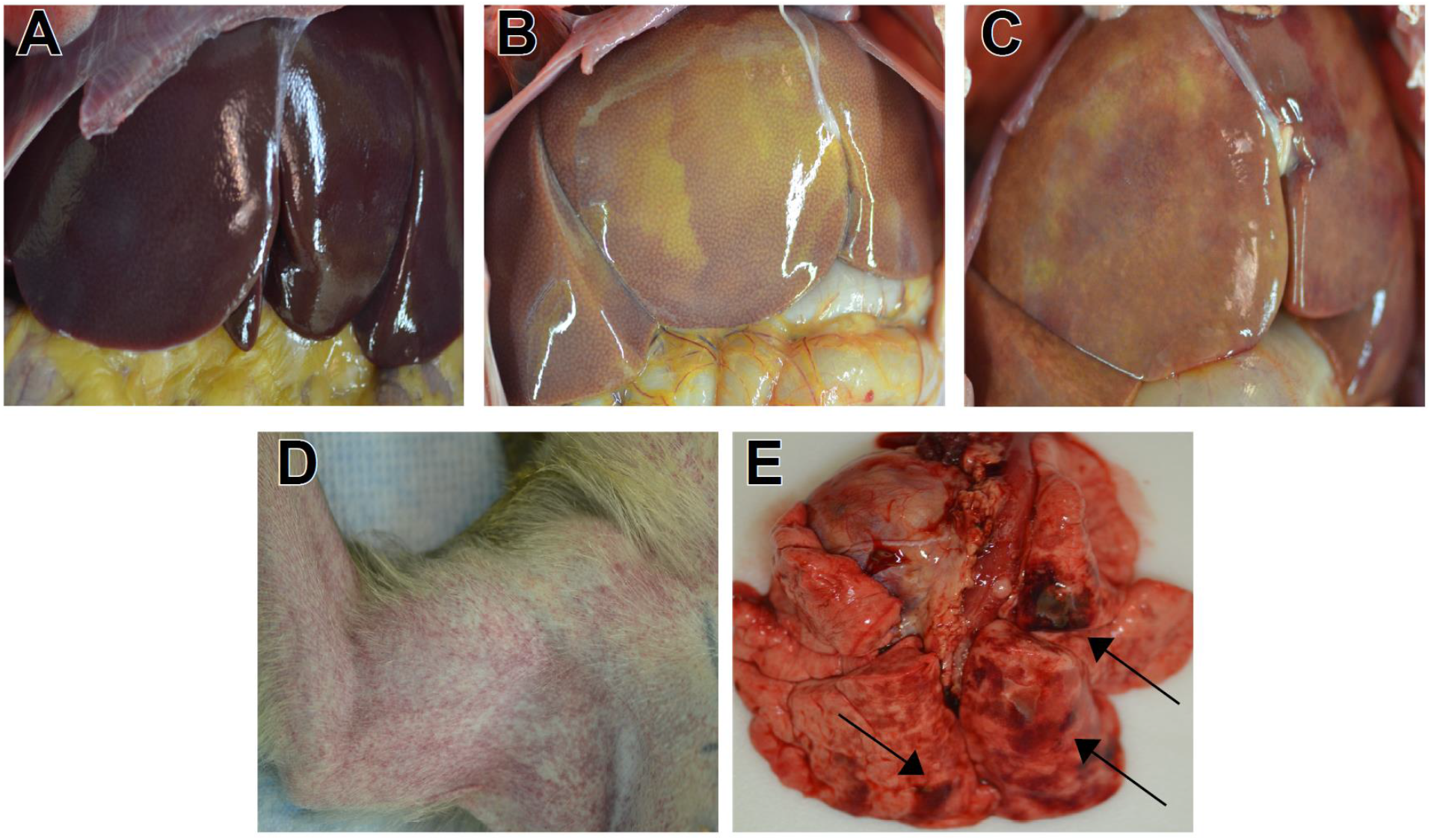
Representative gross pathology in cynomolgus macaques infected with ZEBOV Makona strain via conjunctival route. (**A**) Lack of significant hepatic lesions in a 100PFU survivor (100-5) (**B**) Marked necrotizing hepatitis (500-4), **(C)** Marked diffuse necrotizing hepatitis (100-4)., (**D**) Marked axillary petechial rash (10K-6)**, (E)** Multifocal hemorrhagic interstitial pneumonia (black arrows) (10K-6).

Macaques that succumbed to EVD displayed the expected terminal stage histologic lesions observed in i.m. and small particle aerosol challenged animals despite different challenge doses of 100 PFU (100-4), 500 PFU (500-4) or 10,000 PFU (all animals) of EBOV. Histologic lesions occurring in all macaques that succumbed to EVD included lymphadenitis (axillary, inguinal, and mandibular lymph nodes); tonsillitis; splenitis with lymphoid depletion and fibrin deposition (**Fig 3G,M,S)**; multifocal necrotizing hepatitis (**Fig 3K,Q,W)**; and interstitial pneumonia (**Fig 3I,O,U)**. Other histologic lesions present in at least one macaque that succumbed to EVD included hemorrhagic interstitial pneumonia (**Fig 3U**), uveitis (100-4), adrenalitis (10K-3), tracheitis and esophagitis (10K-2, 10K-3, 10K-4), myocarditis (10K-2, 10K-3), and gastritis (10K-2). Immunolabeling for anti-EBOV antigen was present in macaques that succumbed to disease in the expected cell types, which included individual to small clusters of mononuclear cells within the subcapsular and medullary spaces of the lymph nodes; medullary spaces of the tonsil; red and white pulp of the spleen (**Fig 3H,N,T**); alveolar septate and frequently alveolar macrophages of all lung lobes (**Fig 3J,P,V and insets**) and hepatic sinusoidal lining cells; Kupffer cells; and rarely individual hepatocytes (**Fig 3L,R,X**). Immunohistochemistry (IHC)-positive mononuclear cells were often noted in lesser numbers in the interstitial tissues of the renal cortex, adrenal gland, salivary gland, pancreas, heart, testis, uterus, and prostate. Additionally, mononuclear cells were IHC positive within the dermis or submucosa of the haired skin, nasal mucosa, conjunctiva, urinary bladder, trachea, esophagus, and gastrointestinal tract. IHC-positive mononuclear cells were found within the ciliary body and the draining angle of the eye, adrenal cortical cells, theca cells of the ovary, and the endothelium of small caliber vessels within the meninges and brain parenchyma (**Fig 4E,H,K**). Punctate, cytoplasmic *in situ* hybridization (ISH) signal for viral RNA was abundantly present in the endothelium of small caliber vessels within the brain (**Fig 4F,I,L**). One survivor from the 100 PFU group (100-6) had mild gliosis with an associated focal cluster of IHC-positive neuronal cells within the brainstem (**Fig 4A,B**). Punctate, cytoplasmic ISH was scarcely present in a neuronal cell in the brainstem (**Fig 4C**). No appreciable lesions or IHC labeling for anti-EBOV antigen was present in all other examined organs of this macaque (**Fig 3A-F)**. No prominent lesions or IHC labeling for anti-EBOV antigen were detected in examined tissues of 4/6 macaques from the 100 PFU cohort (100-1,100-2,100-3, 100-5) and 5 of 6 macaques from 500 PFU cohort (500-1, 500-2, 500-3, 100-5, 500-6) (data not shown).

**Figure 3:**
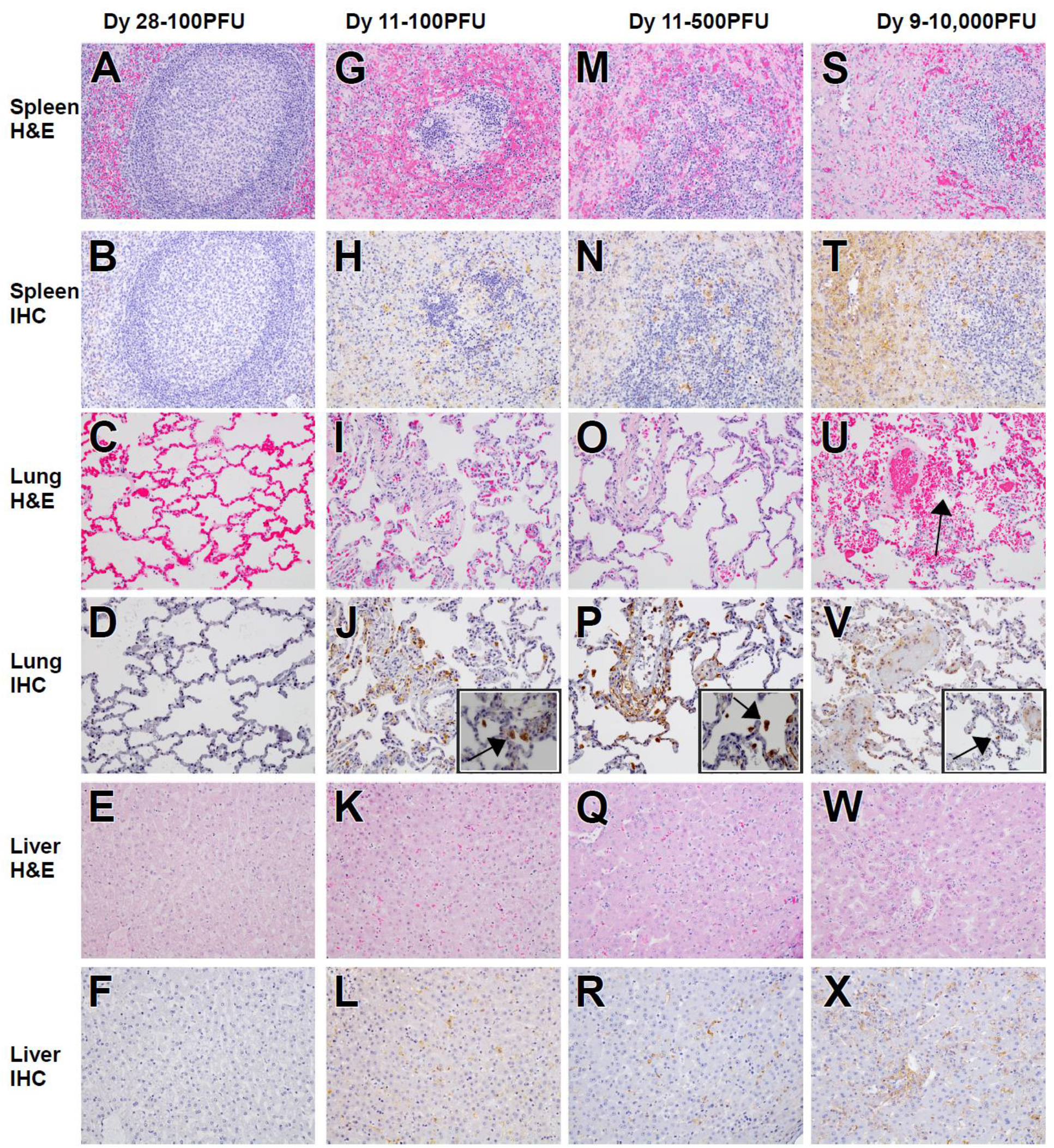
Representative histologic lesions in cynomolgus macaques infected with ZEBOV Makona strain via conjunctival route. Representative tissues of cynomolgus macaques from 100 PFU, survivor (100-6) (**A-F**) and succumbed 11 dpi (100-4) (**G-L**), 500 PFU, succumbed 11 dpi (500-4) (M-R) and 10,000 PFU, succumbed 9 dpi (10K-6) (**S-X**). All images captured at 20x, insets captured at 40x. Hematoxylin and eosin (H&E) staining (**A,G,M,S,C,I,O,U,E,K,Q,W**) and immunohistochemistry (IHC) for anti-EBOV antigen (**B,H,N,T,D,J,J,inset, P,P inset,V, V inset, F, L, R, X**). No significant lesions (NSL) and no significant immunolabeling (NSI) for spleen (**A & B**), lung (**C & D**) and liver (**E & F**) of 100 PFU survivor. Splenitis with lymphoid depletion and fibrin deposition in NHPs that succumb at 100 PFU (**G**), 500 PFU (**M**) and 10,000 PFU (**S**). Diffuse cytoplasmic immunolabeling of mononuclear cells (brown) in red and white pulp of the spleen in those that succumb (**H,N,&T**). Diffuse interstitial pneumonia (**I, O & U**) with alveolar hemorrhage (**U, black arrow**). Diffuse cytoplasmic immunolabeling (brown) of mononuclear cells within the lung (**J, P, & V**) and alveolar spaces (**insets J, P, V, black arrows**). Necrotizing hepatitis (**K, Q, & W**). Diffuse cytoplasmic immunolabeling (brown) of Kupffer cells, sinusoidal lining cells and rarely hepatocytes (**L, R, & X**).

**Figure 4:**
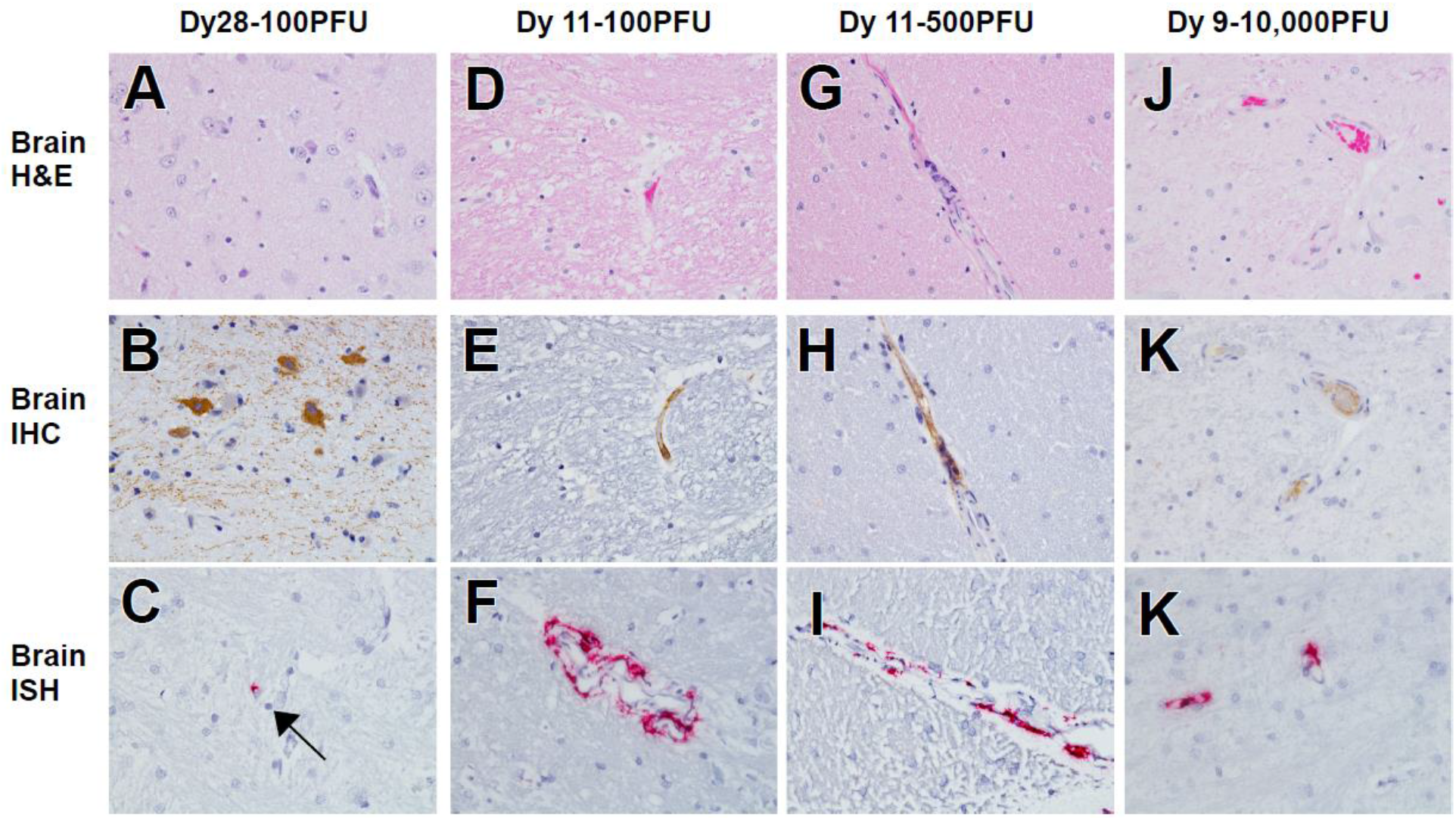
Representative histologic lesions in the brain of cynomolgus macaques infected with ZEBOV Makona strain via conjunctival route. Representative tissues of cynomolgus macaques from 100 PFU, survivor (100-6) (**A-C**) and succumbed 11 dpi (100-4) (**D-F**), 500 PFU, succumbed 11 dpi (500-4) (**G-I**) and 10,000PFU, succumbed 9 dpi (10K-6) (**J-L**). All images captured at 40x. Hematoxylin and eosin (H&E) staining (**A,D,G&J**), immunohistochemistry (IHC) for anti-EBOV antigen (**B,E,H&K**), in situ hybridization (**C,F,I&L**). Mild gliosis (**A,D,G&J**). Diffuse cytoplasmic immunolabeling for anti-EBOV antigen in neuronal cells (B, brown), endothelium of small caliber vessels (**E,H,&K**). Punctate labeling (red) in a single neuronal cell of 100 PFU survivor (100-6) (c, black arrow) and endothelium of small caliber vessels (**F,I,&L**).

### Enzyme-linked immunosorbent assay (ELISA) detection of anti-EBOV antibodies

Terminal sera from all animals surviving to the study endpoint (28 dpi) was analyzed by ELISA and PRNT_50_ for the presence of anti-EBOV antibodies and neutralizing capacity, respectively. In the 100 PFU-challenged cohort, 3/5 animals had detectable titers of anti-EBOV antibodies when assayed against either inactivated virus or GP antigen alone (**Figure S5A**). Titers against inactivated virus were below the threshold for detection in all animals from the 500 PFU-challenged cohort; however, GP-specific IgG was detected in 2/6 animals from this group (**Figure S5B**). Terminal sera from all surviving animals had little to no neutralizing activity against live virus, with none reaching the 50% plaque-reduction threshold for the assay (data not shown).

### Circulating cytokine/chemokine profiling

We assessed levels of select circulating cytokines and chemokines in sera from macaques in the current study and compared them to those from a previous serial euthanasia study utilizing cynomolgus macaques inoculated i.m. with the identical isolate and seed stock of virus (Versteeg, Menicucci et al. 2017). Animals in the current study were sampled daily up to 6 dpi, providing a means of direct comparison of analyte levels during the acute phase of the disease course. For most analytes, patterns of secretion were similar between inoculation routes (**Figure 5**). Notable differences in analyte levels were observed for IL-4 and IL-13, with IL-13 being elevated in macaques inoculated via the conjunctiva versus those inoculated via the i.m. route (**Figure 5I,J**). Moreover, higher levels of IL-13 were observed for macaques challenged via the conjunctiva that succumbed to disease compared to those that survived (**Figure 5J**).

**Figure 5:**
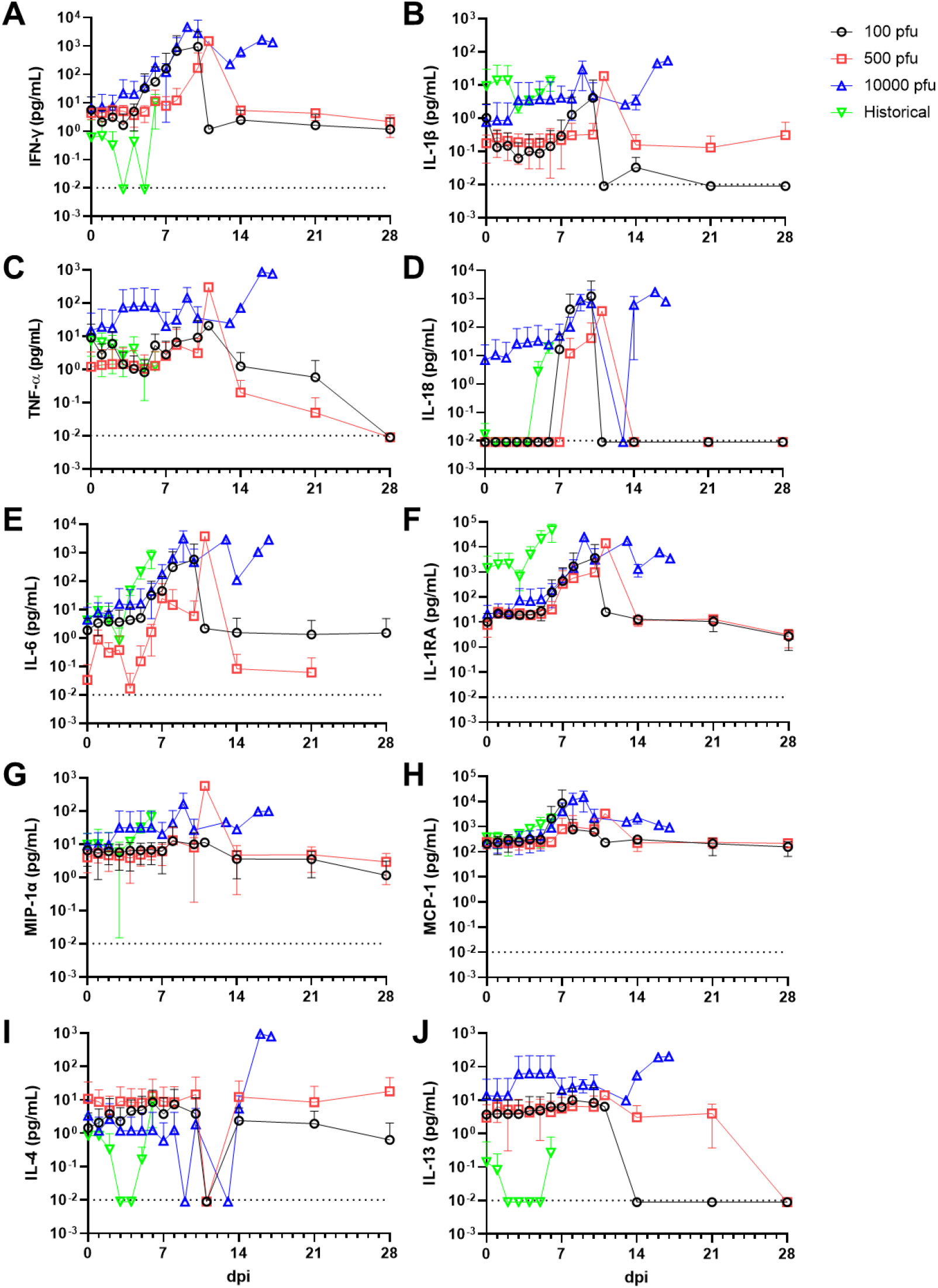
Circulating levels of inflammatory markers. Absolute values of each analyte measured for each subject at the indicated timepoints. Shown is the average value for all animals in the indicated group. Data from historical control animals inoculated i.m. with the homologous virus (Versteeg, Menicucci et al. 2017) were included for statistical purposes. **(A)** IFN-γ; **(B)** IL-1β; **(C)** TNF-α; **(D)** IL-18; **(E)** IL-6; **(F)** IL-1RA; **(G)** MIP-1α; **(H)** MCP-1; **(I)** IL-4; **(J)** IL-13. All assays were run in duplicate reactions.

## Discussion

The development of animal models that recapitulate human diseases is critical to furthering our understanding of the underlying pathological processes and for advancing medical countermeasures. NHPs have proven to be an invaluable resource in accurately modeling the clinical progression of disease, pathology, and severity of EVD in humans (Geisbert, Strong et al. 2015). While sexual transmission of both EBOV and MARV has been well documented, transmission of filoviruses is largely understood to involve exposure to infected animals or via person-to-person through contact with body fluids (e.g., blood, vomitus, saliva), excreta, and possibly fomites; incidental transmission through the facial mucosa is further facilitated through the natural inclination to habitually touch one’s own face, perhaps hundreds of times per day (Suarez and Gallup 1986, Nicas and Best 2008, Kwok, Gralton et al. 2015).

Here, we designed a study with cynomolgus macaques utilizing low to moderate (100 and 500 PFU) and high (10,000 PFU) doses of EBOV variant Makona inoculated via the conjunctiva. A key observation from these studies is that challenge of macaques via the conjunctival route clearly exhibited dose-dependent lethality with a 10,000 PFU dose resulting in uniform lethality, whereas doses of 100 and 500 PFU caused lethal disease in only 1/6 macaques in each cohort. Conversely, the duration and severity of disease in fatal cases, as well as the onset and magnitude of viremia, was not correlated with challenge dose. Surprisingly, while vRNA was detected in some or all tissues from infected subjects regardless of clinical outcome, circulating infectious EBOV was only detected in animals that succumbed to lethal disease. vRNA was found in immunologically privileged potential reservoir sites (eyes and/or gonads) from four surviving animals from the 100 and 500 PFU challenge groups; however, infectious virus was not recovered from any tissues in surviving animals. Detection of circulating vRNA in whole blood was restricted to 3/5 and 1/5 animals from the 100 and 500 PFU challenge cohorts, respectively. While some studies have demonstrated that low doses (0.01-50 PFU) of EBOV or MARV delivered i.m. are sufficient to produce lethal disease in NHPs (Sullivan, Geisbert et al. 2003, Alfson, Avena et al. 2015, Alfson, Avena et al. 2018, Woolsey, Geisbert et al. 2018), the work presented here suggest a higher threshold for productive infection and disease may be necessary for other mucocutaneous of infection with EBOV. Indeed, both rhesus and cynomolgus macaques succumb to disease between 7-8 dpi by i.m. challenge with the Makona isolate of EBOV using a conventional dose of 1000 PFU (Pettitt, Zeitlin et al. 2013, Marzi, Feldmann et al. 2015, Thi, Mire et al. 2015). Comparatively, Jaxx et. al. showed a time to death in rhesus macaques between 7-8 dpi after oral or conjunctival challenge with ~ 100,000 PFU of the Mayinga isolate of EBOV (Jaax, Davis et al. 1996). This contrasts to the findings in our study where the disease onset in fatal cases was considerably delayed (9-17 dpi), although the different viral isolate and species of macaque may influence challenge route-dependent differences. A subsequent exploratory study utilizing small cohorts of cynomolgus macaques demonstrated that low doses (10 or 100 PFU) of EBOV (Makona) delivered via the conjunctiva were non-lethal and only the 100 PFU dose demonstrated low-level viremia as well as seroconversion to the EBOV glycoprotein (Mire, Geisbert et al. 2016). In our study, seroconversion in surviving animals was not uniformly observed across challenged subjects and the neutralizing activity of sera was weak to nonexistent.

Taken together, our studies along with previous studies exploring low-dose EBOV mucosal exposure suggest a substantial difference in threshold for the development of lethal EVD in comparison to i.m. or small particle aerosol exposure. Different exposure routes present both advantages and challenges to productive infection, including differences in the physical and immunological interface. The conjunctiva is one of the most immunologically active mucosal tissues of the external eye (Bielory 2000, Bolanos-Jimenez, Navas et al. 2015). While macrophages, neutrophilic granulocytes, mast cells, and lymphocytes of various lineages are known to inhabit the conjunctiva, the constitutive secretion of immunoglobulin A (IgA) provides additional barriers which may impede productive infection at lower doses. Polymeric/secretory IgA (pIgA/SIgA) can bind some viruses and bacteria leading to their recognition or neutralization by the immune system with varying specificity (Corthesy 2013). Additionally, tear film has anti-microbial properties due to the presence of lysozymes, lactoferrins, lipocalin, and beta-lysine, which can facilitate pathogen defenses including bacterial cell wall lysis, prevention of bacterial and viral binding, inflammation, and detoxification (Bolanos-Jimenez, Navas et al. 2015). IgA has also been found in higher concentrations in tear film than serum (Coyle and Sibony 1986).

We observed increased levels of circulating IL-4 and IL-13 in animals that succumbed to lethal disease, compared to relatively low levels of IL-13 in animals that survived infection. IL-13 is a profibrotic cytokine secreted by type 2 T-helper cells (Th2), mast cells, and basophils (Saw, Offiah et al. 2009), and is involved in goblet cell homeostasis in the respiratory, gastrointestinal, and conjunctival mucosa (De Paiva, Raince et al. 2011). With regards to EBOV, *in vitro* polarization of macrophages to the M2 wound-healing subtype by combined IL-4/IL-13 administration promoted infection by a recombinant vesicular stomatitis virus (rVSV) expressing EBOV GP (rVSV/EBOV GP), but not the parental rVSV vector (Rogers, Brunton et al. 2019). Similarly, *in vivo* treatment or *ex vivo* treatment and implantation of macrophages with IL-4/IL-13 increased disease severity and mortality in mice challenged with rVSV/EBOV GP (Rogers, Brunton et al. 2019). While the IL-13 and IL-4 we measured was systemic, and not localized to the ocular interface, the patterns of expression we observed in this study differed from those observed during the acute phase of infection in i.m.-inoculated macaques (Versteeg, Menicucci et al. 2017), indicating the possibility of a unique role of these cytokines in EBOV pathogenesis via the conjunctival or other mucosal portals. Immunological skewing towards a Th2 phenotype has also been observed to play a role in lethality to MARV infection (Woolsey, Jankeel et al. 2020).

The recent re-emergence of EBOV in Guinea six years after the end of the West African epidemic underpins the importance of understanding pathogenesis and mechanisms of viral clearance as the index case was determined to most likely be recrudescence event from an otherwise recovered patient (Keita, Koundouno et al. 2021). Modeling the complexities associated with survival to filovirus disease in NHPs does afford unique opportunities, but is not without challenges, central of which is the enormous effort and lack of space to conduct long-term studies in high containment facilities. Nonetheless, the value of NHP survivor studies from natural infection or therapeutic studies, particularly when treatment is not initiated until after advanced disease, cannot be understated. Recently, a number of studies with survivors from therapeutic EBOV studies in NHPs demonstrated evidence that replicating virus may still be present in sanctuary sites which are largely immune privileged sites throughout the body, that may still allow for shedding of infectious virus despite potent circulating cellular and humoral immunity (Zeng, Blancett et al. 2017). These sites include reproductive, ocular, and central nervous systems (CNS), all of which evidence has been observed in humans (Mate, Kugelman et al. 2015, Billioux, Smith et al. 2016, Shantha, Yeh et al. 2016). In this study, we provide evidence of replicating virus in the CNS tissues in the context of survival from uninterrupted natural infection.

With a few exceptions, much of the preclinical work for medical countermeasures, including ERVEBO, has utilized the i.m. challenge model (Mire and Geisbert 2017). While certainly important by representing the high-risk scenario of a needle-stick in the hospital or laboratory setting, it is not truly representative of transmission during a natural outbreak, which more likely involves exposure of mucosal surfaces to infected body fluids or excreta. This becomes particularly important in the context of post-exposure treatments. The therapeutic window of i.m. versus mucosal challenge with filoviruses is not equivalent

While some descriptions of survivor models of EBOV infection in the context of treatment have been described, no systematic-in depth-natural history studies exist using a route more likely to be encountered during an outbreak. By using different virus doses via the conjunctival route, we provide details of a novel mucosal challenge model that can be used to interrogate survivor versus lethal EVD signatures, evaluate medical countermeasures, and investigate viral latency.

## Acknowledgments

The authors would like to thank the UTMB Animal Resource Center for husbandry support of laboratory animals and Chad Mire for assistance with the animal studies. Opinions, interpretations, conclusions, and recommendations are those of the authors and are not necessarily endorsed by the University of Texas Medical Branch.

## Author contributions

RWC and TWG conceived and designed the animal challenge experiments. RWC, DJD, JBG, and TWG performed the animal procedures and conducted clinical observations. KNA and VB performed the clinical pathology. KNA performed the PCR and cytokine/chemokine assays. JBG performed the EBOV infectivity assays. CW performed the ELISAs and neutralization assays. NSD performed the IHC and ISH assays and developed the ISH assay. KAF performed gross pathologic, histologic, and immunohistochemical analysis of the data. All authors analyzed the data. RWC, ANP, KF, and TWG wrote the paper. CW edited the paper. All authors had access to the data and approved the final version of the manuscript.

---

**Table 1.**
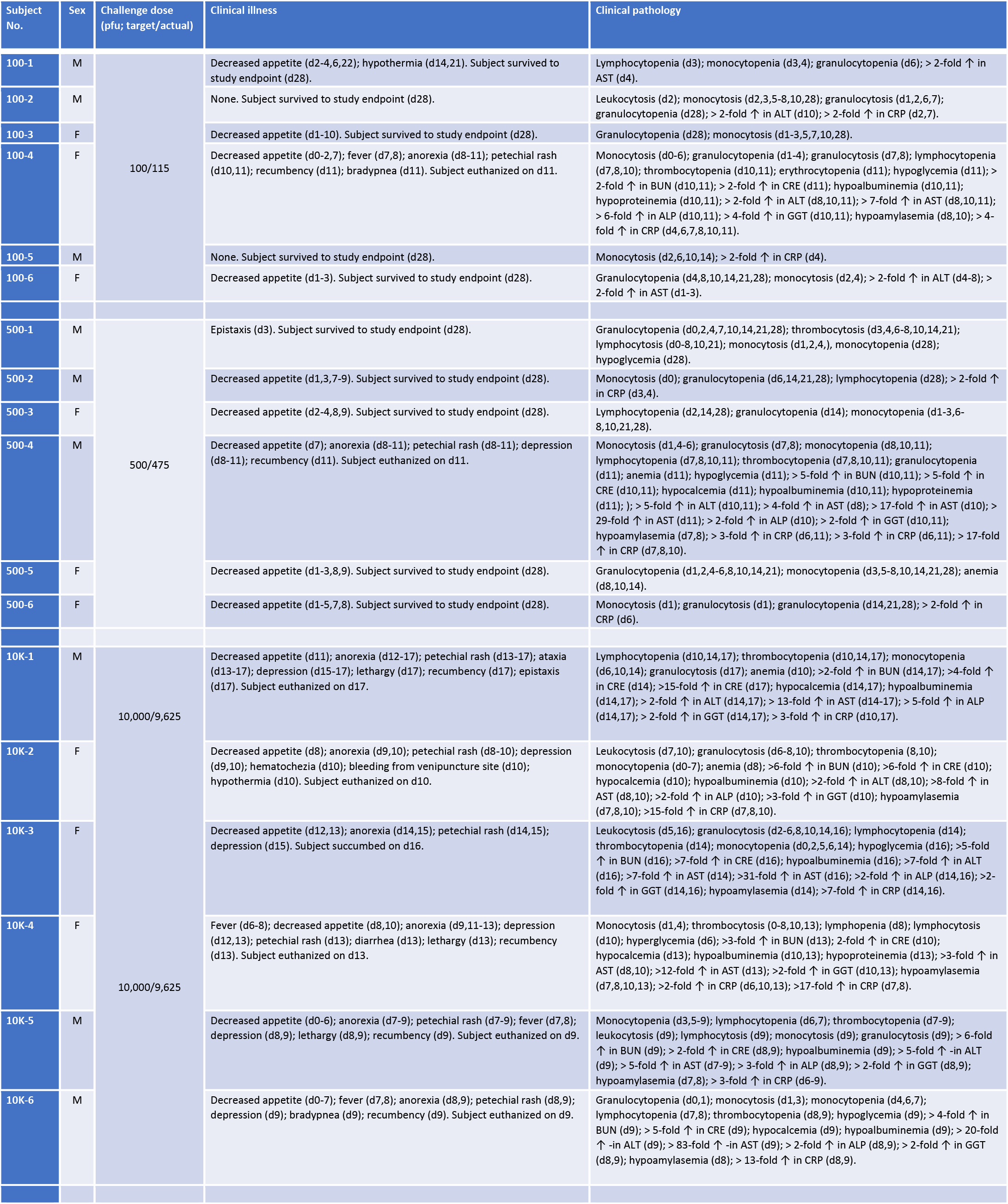
Clinical description and outcome of EBOV challenged cynomolgus macaques. Days after EBOV challenge are in parentheses. All parameters are reported in relation to baseline values (8 days prior to infection; d −8). Lymphopenia, granulopenia, monocytopenia, erythrocytopenia, and thrombocytopenia are defined by a ≥35% drop in numbers of lymphocytes, granulocytes, monocytes, and platelets from baseline, respectively. Leukocytosis, monocytosis and granulocytosis are defined by a two-fold or greater increase in numbers of white blood cells over base line. Thrombocytosis is defined by a two-fold or greater increase in numbers of platelets over baseline. Anemia is defined as a ≥35% decrease in red blood cells, hemoglobin, and hematocrit volume from baseline. Fever is defined as a temperature more than 2.5 °F over baseline, or at least 1.5 °F over baseline and ≥ 103.5 °F. Hypothermia is defined as a temperature ≤3.5°F below baseline. Hyperglycemia is defined as a two-fold or greater increase in levels of glucose. Hypoglycemia is defined by a ≥25% decrease in levels of glucose. Hypocalcemia is defined by a ≥25% decrease in levels of serum calcium. Hypoalbuminemia is defined by a ≥25% decrease in levels of albumin. Hypoproteinemia is defined by a ≥25% decrease in levels of total protein. Hypoamylasemia is defined by a ≥25% decrease in levels of serum amylase. (ALT) alanine aminotransferase, (AST) aspartate aminotransferase, (ALP) alkaline phosphatase, (CRE) Creatinine, (CRP) C-reactive protein, (Hct) hematocrit, (Hgb) hemoglobin

| Subject No. | Sex | Challenge dose (pfu; target/actual) | Clinical illness | Clinical pathology |
| --- | --- | --- | --- | --- |
| 100-1 | M | 100/115 | Decreased appetite (d2-4,6,22); hypothermia (d14,21). Subject survived to study endpoint (d28). | Lymphocytopenia (d3); monocytopenia (d3,4); granulocytopenia (d6); > 2-fold ↑ in AST (d4). |
| 100-2 | M |  | None. Subject survived to study endpoint (d28). | Leukocytosis (d2); monocytosis (d2,3,5-8,10,28); granulocytosis (d1,2,6,7); granulocytopenia (d28); > 2-fold ↑ in ALT (d10); > 2-fold ↑ in CRP (d2,7). |
| 100-3 | F |  | Decreased appetite (d1-10). Subject survived to study endpoint (d28). | Granulocytopenia (d28); monocytosis (d1-3,5,7,10,28). |
| 100-4 | F |  | Decreased appetite (d0-2,7); fever (d7,8); anorexia (d8-11); petechial rash (d10,11); recumbency (d11); bradypnea (d11). Subject euthanized on d11. | Monocytosis (d0-6); granulocytopenia (d1-4); granulocytosis (d7,8); lymphocytopenia (d7,8,10); thrombocytopenia (d10,11); erythrocytopenia (d11); hypoglycemia (d11); > 2-fold ↑ in BUN (d10,11); > 2-fold ↑ in CRE (d11); hypoalbuminemia (d10,11); hypoproteinemia (d10,11); > 2-fold ↑ in ALT (d8,10,11); > 7-fold ↑ in AST (d8,10,11); > 6-fold ↑ in ALP (d10,11); > 4-fold ↑ in GGT (d10,11); hypoamylasemia (d8,10); > 4-fold ↑ in CRP (d4,6,7,8,10,11). |
| 100-5 | M |  | None. Subject survived to study endpoint (d28). | Monocytosis (d2,6,10,14); > 2-fold ↑ in CRP (d4). |
| 100-6 | F |  | Decreased appetite (d1-3). Subject survived to study endpoint (d28). | Granulocytopenia (d4,8,10,14,21,28); monocytosis (d2,4); > 2-fold ↑ in ALT (d4-8); > 2-fold ↑ in AST (d1-3). |
| 500-1 | M | 500/475 | Epistaxis (d3). Subject survived to study endpoint (d28). | Granulocytopenia (d0,2,4,7,10,14,21,28); thrombocytosis (d3,4,6-8,10,14,21); lymphocytosis (d0-8,10,21); monocytosis (d1,2,4,); monocytopenia (d28); hypoglycemia (d28). |
| 500-2 | M |  | Decreased appetite (d1,3,7-9). Subject survived to study endpoint (d28). | Monocytosis (d0); granulocytopenia (d6,14,21,28); lymphocytopenia (d28); > 2-fold ↑ in CRP (d3,4). |
| 500-3 | F |  | Decreased appetite (d2-4,8,9). Subject survived to study endpoint (d28). | Lymphocytopenia (d2,14,28); granulocytopenia (d14); monocytopenia (d1-3,6-8,10,21,28). |
| 500-4 | M |  | Decreased appetite (d7); anorexia (d8-11); petechial rash (d8-11); depression (d8-11); recumbency (d11). Subject euthanized on d11. | Monocytosis (d1,4-6); granulocytosis (d7,8); monocytopenia (d8,10,11); lymphocytopenia (d7,8,10,11); thrombocytopenia (d7,8,10,11); granulocytopenia (d11); anemia (d11); hypoglycemia (d11); > 5-fold ↑ in BUN (d10,11); > 5-fold ↑ in CRE (d10,11); hypocalcemia (d11); hypoalbuminemia (d10,11); hypoproteinemia (d11); > 5-fold ↑ in ALT (d10,11); > 4-fold ↑ in AST (d8); > 17-fold ↑ in AST (d10); > 29-fold ↑ in AST (d11); > 2-fold ↑ in ALP (d10); > 2-fold ↑ in GGT (d10,11); hypoamylasemia (d7,8); > 3-fold ↑ in CRP (d6,11); > 3-fold ↑ in CRP (d6,11); > 17-fold ↑ in CRP (d7,8,10). |
| 500-5 | F |  | Decreased appetite (d1-3,8,9). Subject survived to study endpoint (d28). | Granulocytopenia (d1,2,4-6,8,10,14,21); monocytopenia (d3,5-8,10,14,21,28); anemia (d8,10,14). |
| 500-6 | F |  | Decreased appetite (d1-5,7,8). Subject survived to study endpoint (d28). | Monocytosis (d1); granulocytosis (d1); granulocytopenia (d14,21,28); > 2-fold ↑ in CRP (d6). |
| 10K-1 | M | 10,000/9,625 | Decreased appetite (d11); anorexia (d12-17); petechial rash (d13-17); ataxia (d13-17); depression (d15-17); lethargy (d17); recumbency (d17); epistaxis (d17). Subject euthanized on d17. | Lymphocytopenia (d10,14,17); thrombocytopenia (d10,14,17); monocytopenia (d6,10,14); granulocytosis (d17); anemia (d10); >2-fold ↑ in BUN (d14,17); >4-fold ↑ in CRE (d14); >15-fold ↑ in CRE (d17); hypocalcemia (d14,17); hypoalbuminemia (d14,17); > 2-fold ↑ in ALT (d14,17); > 13-fold ↑ in AST (d14-17); > 5-fold ↑ in ALP (d14,17); > 2-fold ↑ in GGT (d14,17); > 3-fold ↑ in CRP (d10,17). |
| 10K-2 | F |  | Decreased appetite (d8); anorexia (d9,10); petechial rash (d8-10); depression (d9,10); hematochezia (d10); bleeding from venipuncture site (d10); hypothermia (d10). Subject euthanized on d10. | Leukocytosis (d7,10); granulocytosis (d6-8,10); thrombocytopenia (8,10); monocytopenia (d0-7); anemia (d8); >6-fold ↑ in BUN (d10); >6-fold ↑ in CRE (d10); hypocalcemia (d10); hypoalbuminemia (d10); >2-fold ↑ in ALT (d8,10); >8-fold ↑ in AST (d8,10); >2-fold ↑ in ALP (d10); >3-fold ↑ in GGT (d10); hypoamylasemia (d7,8,10); >15-fold ↑ in CRP (d7,8,10). |
| 10K-3 | F |  | Decreased appetite (d12,13); anorexia (d14,15); petechial rash (d14,15); depression (d15). Subject succumbed on d16. | Leukocytosis (d5,16); granulocytosis (d2-6,8,10,14,16); lymphocytopenia (d14); thrombocytopenia (d14); monocytopenia (d0,2,5,6,14); hypoglycemia (d16); >5-fold ↑ in BUN (d16); >7-fold ↑ in CRE (d16); hypoalbuminemia (d16); >7-fold ↑ in ALT (d16); >7-fold ↑ in AST (d14); >31-fold ↑ in AST (d16); >2-fold ↑ in ALP (d14,16); >2-fold ↑ in GGT (d14,16); hypoamylasemia (d14); >7-fold ↑ in CRP (d14,16). |
| 10K-4 | F | 10,000/9,625 | Fever (d6-8); decreased appetite (d8,10); anorexia (d9,11-13); depression (d12,13); petechial rash (d13); diarrhea (d13); lethargy (d13); recumbency (d13). Subject euthanized on d13. | Monocytosis (d1,4); thrombocytosis (0-8,10,13); lymphopenia (d8); lymphocytosis (d10); hyperglycemia (d6); >3-fold ↑ in BUN (d13); 2-fold ↑ in CRE (d10); hypocalcemia (d13); hypoalbuminemia (d10,13); hypoproteinemia (d13); >3-fold ↑ in AST (d8,10); >12-fold ↑ in AST (d13); >2-fold ↑ in GGT (d10,13); hypoamylasemia (d7,8,10,13); >2-fold ↑ in CRP (d6,10,13); >17-fold ↑ in CRP (d7,8). |
| 10K-5 | M |  | Decreased appetite (d0-6); anorexia (d7-9); petechial rash (d7-9); fever (d7,8); depression (d8,9); lethargy (d8,9); recumbency (d9). Subject euthanized on d9. | Monocytopenia (d3,5-9); lymphocytopenia (d6,7); thrombocytopenia (d7-9); leukocytosis (d9); lymphocytosis (d9); monocytosis (d9); granulocytosis (d9); > 6-fold ↑ in BUN (d9); > 2-fold ↑ in CRE (d8,9); hypoalbuminemia (d9); > 5-fold ↑ in ALT (d9); > 5-fold ↑ in AST (d7-9); > 3-fold ↑ in ALP (d8,9); > 2-fold ↑ in GGT (d8,9); hypoamylasemia (d7,8); > 3-fold ↑ in CRP (d6-9). |
| 10K-6 | M |  | Decreased appetite (d0-7); fever (d7,8); anorexia (d8,9); petechial rash (d8,9); depression (d9); bradypnea (d9); recumbency (d9). Subject euthanized on d9. | Granulocytopenia (d0,1); monocytosis (d1,3); monocytopenia (d4,6,7); lymphocytopenia (d7,8); thrombocytopenia (d8,9); hypoglycemia (d9); > 4-fold ↑ in BUN (d9); > 5-fold ↑ in CRE (d9); hypocalcemia (d9); hypoalbuminemia (d9); > 20-fold ↑ in ALT (d9); > 83-fold ↑ in AST (d9); > 2-fold ↑ in ALP (d8,9); > 2-fold ↑ in GGT (d8,9); hypoamylasemia (d8); > 13-fold ↑ in CRP (d8,9). |

**Fig S1:**
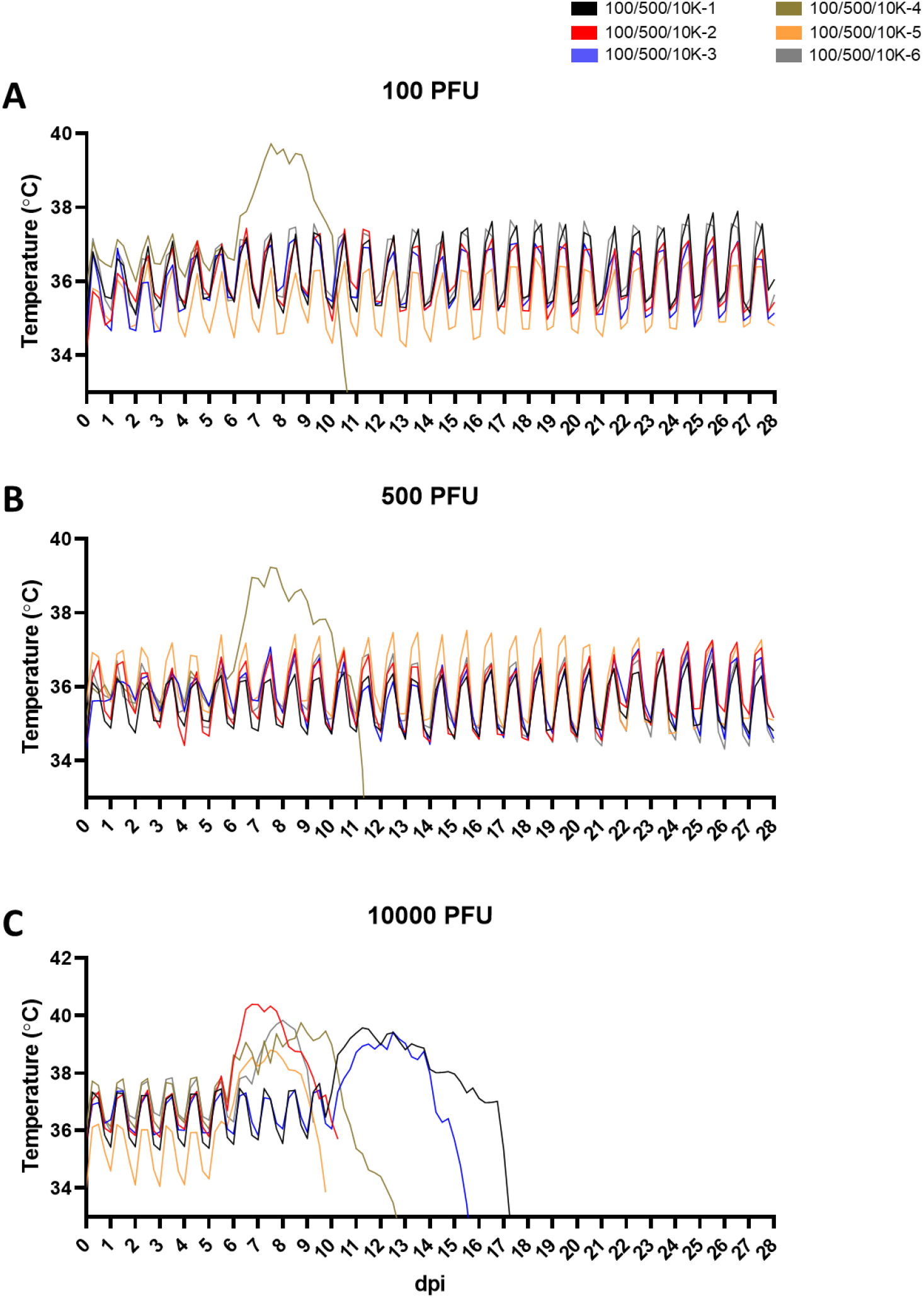
Core body temperature telemetry of EBOV-challenged cynomolgus macaques. Core body temperatures from each macaque were measured in 15-minute increments throughout the study duration via intraperitoneal implantation of telemetric temperature loggers (described in “Methods”). Each data point represents the 6-hour rolling average (i.e., average of 24 measurements within a 6 hour block) for each animal. **(A)** Core temperature measurements for the 100 PFU challenge cohort; **(B)** Core temperature measurements for the 500 PFU challenge cohort; **(C)** Core temperature measurements for the 10,000 PFU challenge cohort.

**Figure S2:**
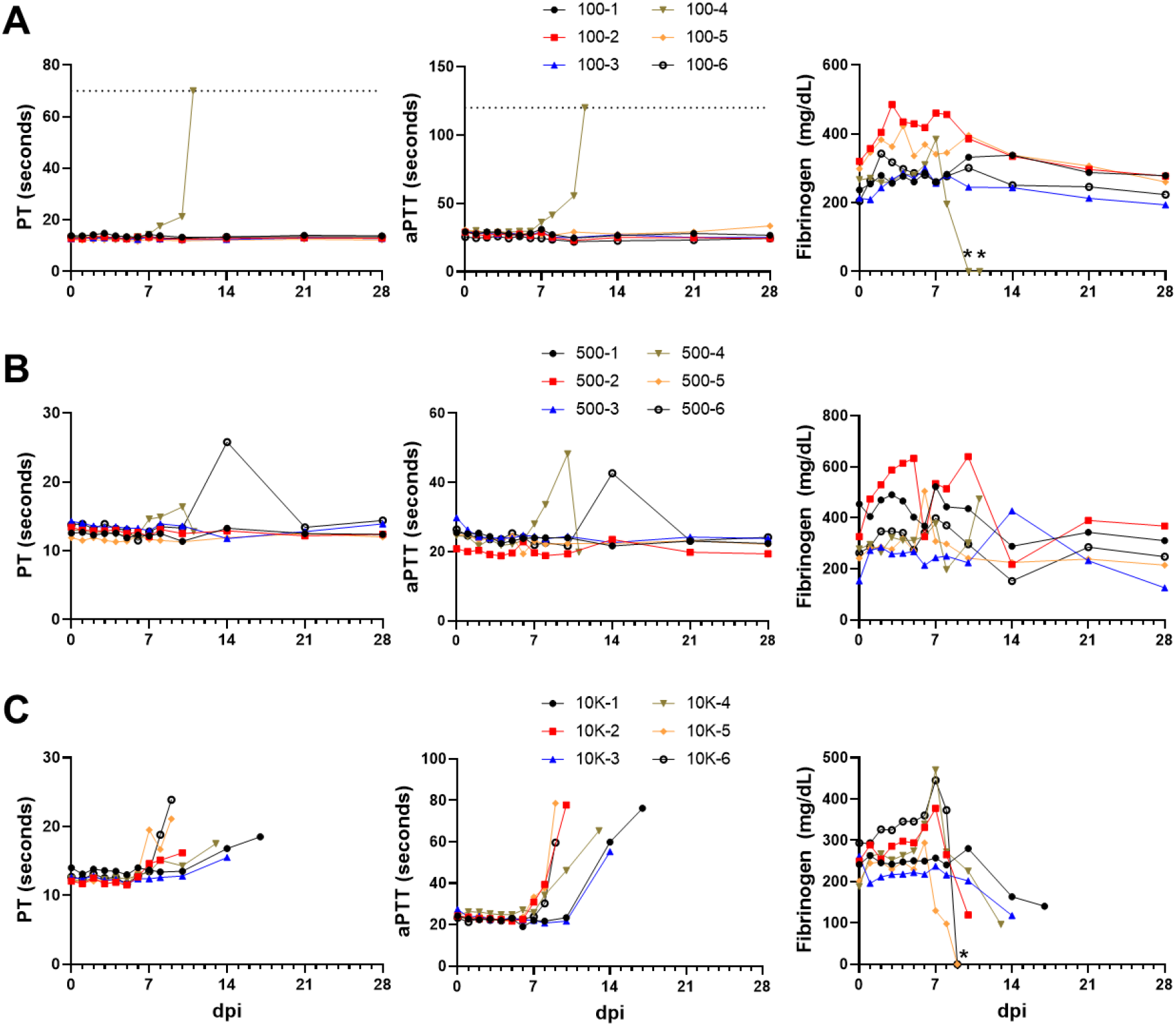
Coagulopathy profiles of cynomolgus macaques challenged with EBOV virus. Measurement of coagulation parameters for cynomolgus macaques challenged with 100 PFU **(A)**, 500 PFU **(B)**, or 10,000 PFU **(C)** EBOV-Makona. For each animal, the prothrombin time (PT), activated partial thromboplastin time (aPTT), and circulating fibrinogen were measured at the indicated timepoints. Horizontal dashed lines in the PT and aPTT panels in **(A)** indicate the upper limit of detection for the assay. Asterisks in the fibrinogen panels in **(A)** and **(C)** indicate undetectable levels.

**Figure S3:**
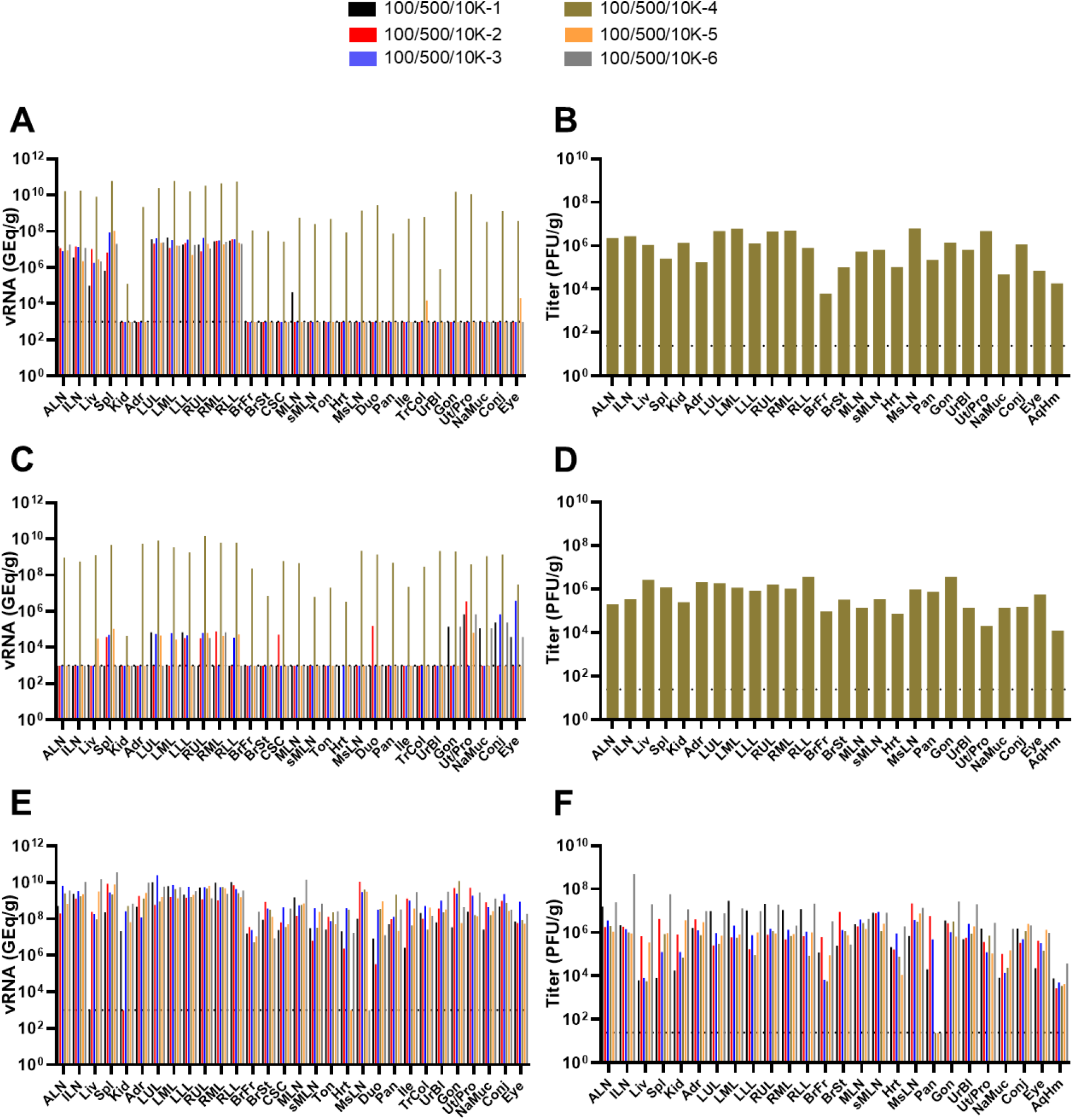
Determination of tissue viral burden in EBOV-challenged macaques. Viral load was determined by RT-qPCR detection of EBOV vRNA **(A,C,E)** and plaque titration (**B,D,E)** from selected tissues harvested at necropsy. For all panels, individual data points represent the mean of two technical replicates. Dashed horizontal lines indicate the limit of detection (LOD) for the assay (1000 GEq/g tissue for RT-qPCR; 25 PFU/g tissue for plaque titration). Values below the LOD for RT-qPCR and plaque assays were plotted as 999 GEq/g and 24 PFU/g tissue, respectively. Missing data indicates the tissue was not collected or assayed for that subject. Abbreviations for tissues: ALN: Axillary lymph node; ILN: inguinal lymph node; Liv: liver; Spl: spleen; Kid: kidney; Adr: adrenal gland; LUL: left upper lung; LML: left middle lung; LLL: left lower lung; RUL: right upper lung; RML: right middle lung; RLL: right lower lung; BrFr: brain frontal cortex; BrSt: brain stem; CSC: cervical spinal cord; MLN: mandibular lymph node; sMLN: submandibular lymph node; Ton: tonsil; Hrt: heart; MsLN: mesenteric lymph node; Duo: duodenum; Pan: pancreas; Ile: ileum; TrCol: transverse colon; UrBl: urinary bladder; Gon: gonad; Ut/Pro: uterus/prostate; NaMuc: nasal mucosa; Conj: conjunctiva.

**Figure S4:**
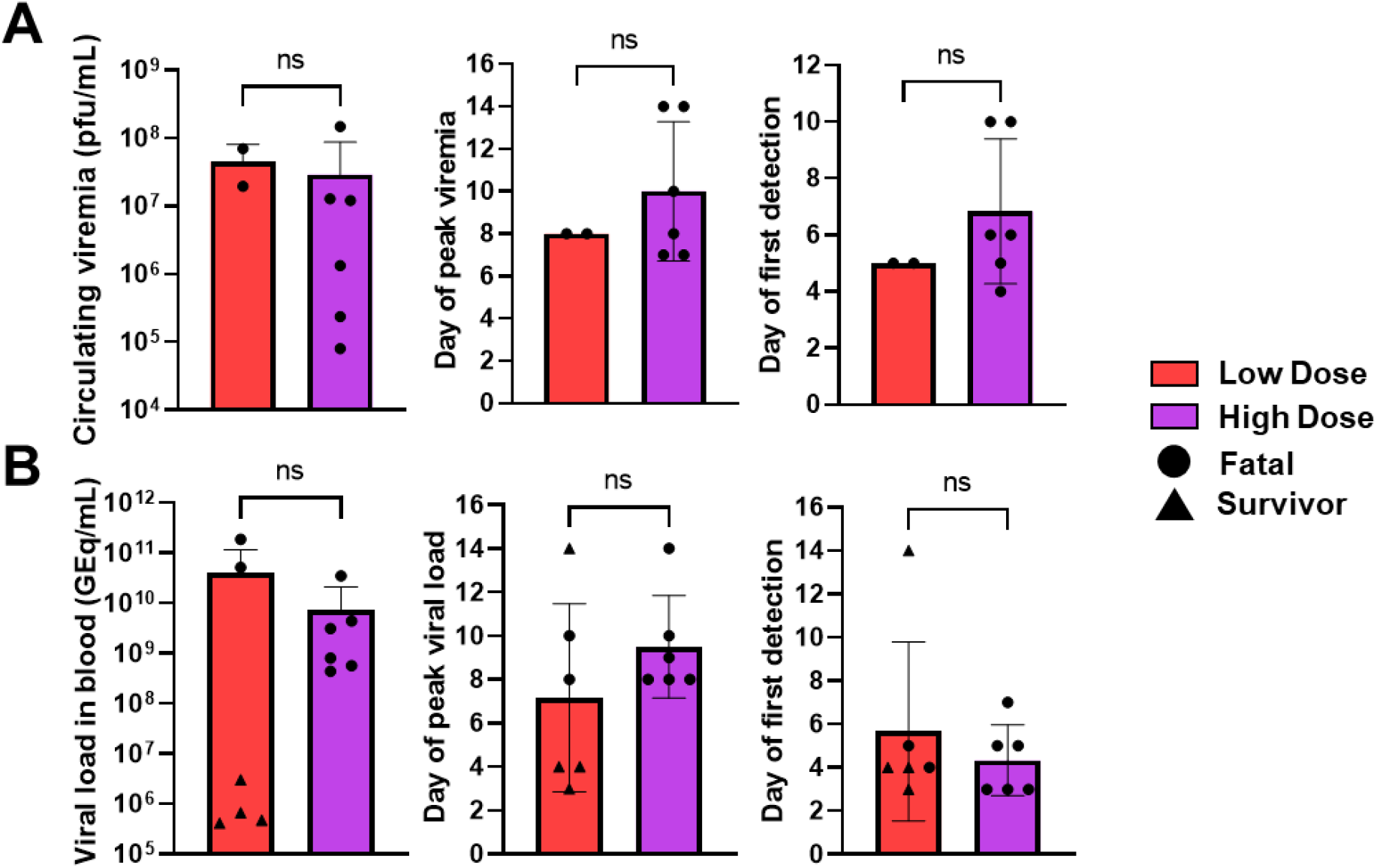
Comparison of viral RNA and infectious virus in whole blood and plasma from EBOV-challenged cynomolgus macaques. The peak viral load (irrespective of which day it was detected), the day peak viral load was detected, and the day infectious virus **(A)** or EBOV vRNA **(B)** was first detected in whole blood or plasma, respectively, was compared between macaques challenged with “low” (100 and 500 PFU) or “high” (10,000 PFU) doses of EBOV. Statistical significance was determined by unpaired t-test with Welch’s correction.

**Fig S5:**
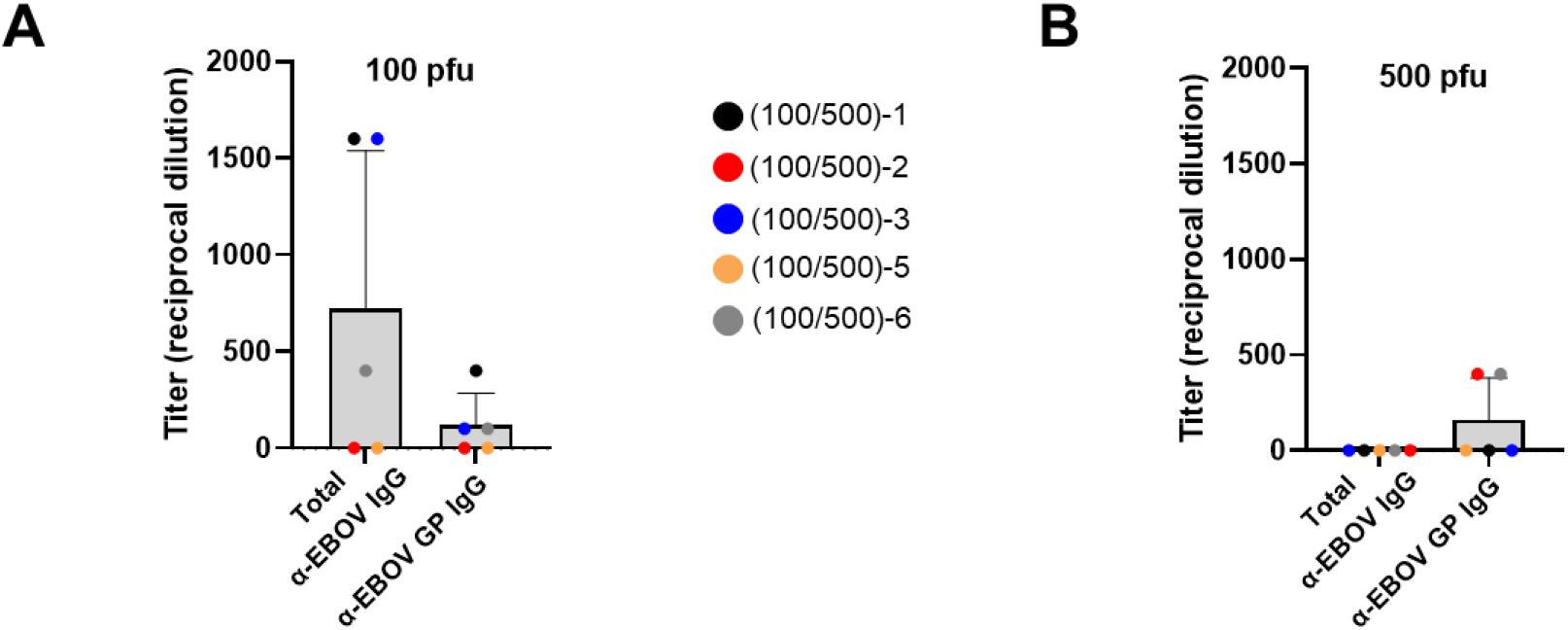
Determination serum IgG titers in EBOV-challenged macaques. ELISA-based quantification of serum anti-IgG titers from survivors at the study endpoint (28 dpi). IgG titers were measured against irradiated virus or GP antigen. Bars represent the mean IgG titer ± SD, values from individual animals are represented as colored circles within the bars.

## Materials and Methods

### Challenge Virus

The EBOV Makona variant seed stock originated from serum from a fatal case during the 2014 outbreak in Guékédou, Guinea (Ebola virus/H.sapiens-wt/GIN/2014/Makona-C07, accession number KJ660347.2) and was passaged twice in authenticated Vero E6 cells obtained from ATCC (ATCC, CRL-1586) (Mire, Matassov et al. 2015, Thi, Mire et al. 2015).

### Animal Challenge

Eighteen healthy adult cynomolgus macaques (*Macaca fascicularis*) of Chinese origin (weight range, 4.3-7.0 kg; age range, 4-8 years; PreLabs) were used for these studies. Three sex-balanced challenge cohorts were established with 6 animals per cohort. For continuous core body temperature measurements, a DST micro-T implantable temperature logger (Star–Oddi, Gardabaer, Iceland) was surgically implanted to each animal prior to study initiation; data recording was set to 10 or 15 minute intervals. Each cohort was exposed to a target dose of 100, 500, or 10,000 PFU of EBOV Makona by droplet administration into the medial canthus of each eye. For each challenge cohort, animals underwent physical examinations and blood specimens were collected at the time of challenge (day 0) and on days 1, 2, 3, 4, 5, 6, 7, 8, 10, 14, 21, and 28 post infection. Animals were monitored daily and scored for disease progression with an internal filovirus scoring protocol approved by the UTMB Institutional Animal Care and Use Committee (IACUC) and the USAMRDC Animal Care and Use Review Office (ACURO). The scoring changes measured from baseline included posture/activity level, attitude/behavior, food and water intake, weight, respiration, and disease manifestations such as visible rash, hemorrhage, ecchymosis, or flushed skin. A score of ≥ 9 indicated that an animal met criteria for euthanasia. This study was not blinded. Any surviving animals were euthanized on day 28. Animal studies were completed under Biosafety Level 4 containment at the Galveston National Laboratory (GNL) and were approved by the University of Texas Medical Branch (UTMB) IACUC, in accordance with state and federal statutes and regulations relating to experiments involving animals, and the UTMB Institutional Biosafety Committee.

### Hematologic and Serum Biochemical Analysis

Total white blood cell counts, white blood cell differentials, red blood cell counts, platelet counts, hematocrit values, total hemoglobin concentrations, mean cell volumes, mean corpuscular volumes, and mean corpuscular hemoglobin concentrations were analyzed in blood specimens collected in tubes containing ethylenediaminetetraacetic acid, using a laser based hematologic analyzer (Beckman Coulter). Serum samples were tested for concentrations of albumin, amylase, alanine aminotransferase, aspartate aminotransferase, alkaline phosphatase, γ glutamyl transferase, glucose, cholesterol, total protein, total bilirubin, blood urea nitrogen, creatinine, and C-reactive protein by using a Piccolo point-of-care analyzer and Biochemistry Panel Plus analyzer discs (Abaxis). Citrated plasma samples were analyzed for coagulation parameters PT, APTT, thrombin time, and fibrinogen on the STart4 instrument using the PTT Automate, STA Neoplastine CI Plus, STA Thrombin, and Fibri-Prest Automate kits, respectively (Diagnostica Stago).

### Detection of Viremia and Viral RNA

RNA was isolated from whole blood utilizing the Viral RNA mini-kit (Qiagen) using 100 μl of blood added to 600 μl of the viral lysis buffer. Primers and probe targeting the VP30 gene of EBOV were used for real-time quantitative PCR (RT-qPCR) with the following probes: EBOV, 6-carboxyfluorescein (FAM)-5’ CCG TCA ATC AAG GAG CGC CTC 3’-6 carboxytetramethylrhodamine (TAMRA) (Life Technologies).Viral RNA was detected using the CFX96 detection system (Bio-Rad Laboratories, Hercules, CA) in one-step probe RT-qPCR kits (Qiagen) with the following cycle conditions: 50°C for 10 min, 95°C for 10 s, and 40 cycles of 95°C for 10 s and 57°C for 30 s. Threshold cycle (CT) values representing viral genomes were analyzed with CFX Manager software, and the data are shown as genome equivalents (GEq) per milliliter. To create the GEq standard, RNA from viral stocks was extracted, and the number of strain-specific genomes was calculated using Avogadro’s number and the molecular weight of each viral genome.

Virus titration was performed for by plaque assay with Vero E6 cells from all plasma samples as previously described (Mire, Matassov et al. 2015, Thi, Mire et al. 2015). Briefly, increasing 10-fold dilutions of the samples were adsorbed to Vero E6 monolayers in duplicate wells (200 μL); the limit of detection was 25 PFU/mL.

### Anti-EBOV GP IgG ELISA

Sera collected at the indicated time points were tested for anti-EBOV immunoglobulin G (IgG) antibodies by standard ELISA. For GP-specific IgG titers, MaxiSorp clear flat-bottom 96-well plates (44204 ThermoFisher, Rochester, NY) were coated overnight with 15 ng/well (0.15mL) of recombinant EBOV Makona GPΔTM (ΔTM: transmembrane region absent; Integrated Biotherapeutics, Gaithersburg, MD) in a sodium carbonate/bicarbonate solution (pH 9.6). Antigen-adsorbed wells were subsequently blocked with 4% bovine serum antigen (BSA) in 1 x PBS for at least two hours. Sera were initially diluted 1:100 and then two-fold through 1:12800 in ELISA diluent (1% BSA in 1× PBS, and 0.2% Tween-20). For total IgG titers, plates were coated with irradiated EBOV-Makona antigen or normal Vero E6 antigen kindly provided by Dr. Thomas W. Ksiazek (UTMB). Sera were initially diluted 1:100 and then four-fold through 1:6400 in 3% BSA in 1× PBS. After a one-hour incubation, plates were washed six times with wash buffer (1 x PBS with 0.2% Tween-20) and incubated for an hour with a 1:2500 dilution of horseradish peroxidase (HRP)-conjugated anti-rhesus IgM or IgG antibody (Fitzgerald Industries International, Acton, MA). RT SigmaFast O-phenylenediamine (OPD) substrate (P9187, Sigma, St. Louis, MO) was added to the wells after six additional washes to develop the colorimetric reaction. The reaction was stopped with 3M sulfuric acid 5-10 minutes after OPD addition and absorbance values were measured at a wavelength of 492nm on a spectrophotometer (Molecular Devices Emax system, Sunnyvale, CA). Absorbance values were normalized by subtracting normal/uncoated wells from antigen-coated wells at the corresponding serum dilution. For total IgG titers, a sum OD value > 0.6 was required to denote a positive sample. End-point titers were defined as the reciprocal of the last adjusted serum dilution with a value ≥ 0.20.

### Histopathologic, Immunohistochemical (IHC), and In Situ Hybridization (ISH) Analyses

Necropsy was performed on all subjects. Tissue samples from major organs were collected for histopathological and IHC examination, immersion fixed in 10% neutral buffered formalin, and processed for histopathologic analysis as previously described (Mire, Matassov et al. 2015, Thi, Mire et al. 2015). For IHC analysis, specific anti-EBOV immunoreactivity was detected using an anti-EBOV VP40 protein rabbit primary antibody (Integrated BioTherapeutics) at a 1:4000 dilution. Tissue sections were processed for IHC analysis, using the Dako Autostainer (Dako). Secondary antibody used was biotinylated goat anti-rabbit IgG (Vector Laboratories) at 1:200 followed by Vector horseradish peroxidase streptavidin, R.T.U (Vector Laboratories) for 30 minutes. Slides were developed with Dako DAB chromagen (Dako) for 5 minutes and counterstained with hematoxylin for 45 seconds. Methods for visualization with red chromogen were identical, except Vector Streptavidin Alkaline Phosphatase (Vector Laboratories) was used at a 1:200 dilution for 20 minutes, and slides were developed with Bio-Red (Biopath Laboratories) for 7 minutes and counterstained with hematoxylin for 45 seconds. EBOV RNA ISH in formalin-fixed paraffin embedded (FFPE) tissues was performed on representative tissue sections of brain using the RNAscope 2.5 high definition (HD) RED kit (Advanced Cell Diagnostics, Newark, CA) according to the manufacturer’s instructions. 20 ZZ probe pairs targeting the genomic EBOV nucleoprotein (NP) gene were designed and synthesized by Advanced cell Diagnostics (catalog 448581). After sectioning, deparaffinization with xylene and graded ethanol washes and peroxidase blocking, the sections were heated in RNAscope Target Retrieval Reagent Buffer (Advanced Cell Diagnostics catalogue 322000) for 45 minutes and then air-dried overnight. The sections were then digested with Protease IV (catalogue 322336) at 40C in the HybEZ oven (HybEZ, Advanced Cell Diagnostics catalogue 321711) for 30 minutes. Sections were exposed to ISH target probe and incubated at 40C in the HybEZ oven for 2 hours. After rinsing, the signal was amplified using the manufacturer provided pre-amplifier and amplifier conjugated to alkaline phosphatase and incubated with a red substrate-chromogen solution for 10 minutes, counterstained with hematoxylin, air-dried, and coverslipped.

### EBOV neutralization assay

Neutralization assays were performed by measuring plaque reduction in a constant virus:serum dilution format as previously described (Jones, Feldmann et al. 2005). Briefly, a standard amount of EBOV strain Makona (~ 100 PFU) was incubated with serial two-fold dilutions of the serum sample for 60 min. The mixture was used to inoculate Vero E6 cells (ATCC, CRL-1586) for 60 min. Cells were overlayed with an agar medium, incubated for 7 days, and plaques were counted 48 h after neutral red staining. Endpoint titers were determined by the dilution of serum which neutralized 50% of the plaques (PRNTT_50_).

### Bead-based multiplex immunoassays

The concentrations of circulating cytokines, chemokines, and other analytes were assayed using bead-based multiplex technology. Irradiated plasma samples were incubated with magnetic beads from Milliplex NHP cytokine premixed 23-plex panel (EMD Millipore, Billerica, MA) kits according to the recommendations provided. Analytes measured included IL-1β, IL-1 receptor antagonist (IL-1RA), IL-2, IL-4, IL-5, IL-6, IL-8, IL-10, IL-12/23 (p40), IL-13, IL-15, IL-17, IL-18, gamma interferon (IFN-γ), granulocyte colony-stimulating factor (G-CSF), granulocyte-macrophage colony-stimulating factor (GM-CSF), monocyte chemoattractant protein 1 (MCP-1), macrophage inflammatory protein 1α (MIP-1α), MIP-1β, tumor necrosis factor alpha (TNF-α), transforming growth factor alpha (TGF-α), soluble CD40 ligand (sCD40L), and vascular endothelial growth factor (VEGF). The concentrations in each plasma sample were measured using a Bioplex-200 array system (Bio-Rad, Hercules, CA).

